# Ecological metabolomics of tropical tree communities across an elevational gradient: Implications for chemically-mediated biotic interactions and species diversity

**DOI:** 10.1101/2023.10.04.560880

**Authors:** David Henderson, Brian E. Sedio, J. Sebastián Tello, Leslie Cayola, Alfredo F. Fuentes, Belen Alvestegui, Nathan Muchhala, Jonathan A. Myers

## Abstract

Seminal hypotheses in ecology and evolution postulate that stronger and more specialized biotic interactions contribute to higher species diversity at lower elevations and latitudes. Plant-chemical defenses mediate biotic interactions between plants and their natural enemies and provide a highly dimensional trait space in which chemically mediated niches may facilitate plant species coexistence. However, the role of chemically mediated biotic interactions in shaping plant communities remains largely untested across large-scale ecological gradients. To test this hypothesis, we used ecological metabolomics to quantify the chemical dissimilarity of foliar metabolomes among 473 tree species (906 unique species-plot combinations) in 16 tropical tree communities along an elevational gradient in Madidi National Park, Bolivia. We predicted that chemical dissimilarity among co-occurring tree species would be greater, and chemical phylogenetic signal lower, in communities with greater tree species richness and warmer, wetter, and less-seasonal climates, as pressure from natural enemies is likely to be greater in these locales. Further, we predicted that these relationships should be especially pronounced for secondary metabolites derived from biosynthetic pathways known to include anti-herbivore and antimicrobial defenses than for primary metabolites. We found that median chemical dissimilarity among tree species with respect to all metabolites and secondary metabolites increased with tree species richness, decreased with elevation, and increased along a principal component of climatic variation that reflected increasing temperature and precipitation and decreasing seasonality. In contrast, median chemical dissimilarity among tree species with respect to primary metabolites was unrelated to tree species richness, elevation, or the principal component of climatic variation. Furthermore, phylogenetic signal of secondary and primary metabolites decreased with tree species richness. Among tree communities in moist forests, phylogenetic signal of secondary metabolites also increased with elevation and decreased with the temperature and precipitation. Our results support the hypothesis that chemically mediated biotic interactions shape elevational diversity gradients by imposing stronger selection for interspecific divergence in plant chemical defenses in warmer, wetter, and more stable climates. Our study also illustrates the promise of ecological metabolomics in the study of biogeography, community ecology, and complex species interactions in high-diversity ecosystems.

## Introduction

Foundational hypotheses in ecology and evolution posit that stronger and more specialized biotic interactions contribute to large-scale gradients in biological diversity (Schemske et al., 2009). Wallace (1878) and Dobzhansky (1950) proposed that biotic interactions comprise a stronger selective force than the abiotic environment in the tropics. However, the mechanisms by which tropical forests may facilitate the ecological coexistence of hundreds to thousands of tree species remain unclear (Wright 2002). Unlike animals, which can exploit distinct resources, nearly all plants require light, water, CO_2_, and a few shared micronutrients, so opportunities for resource-based niche differentiation are few (Hubbell 2001). In contrast to resource-based niche axes, the nearly infinite variety of insect herbivores and microbial pathogens provides a highly multidimensional space within which plant species can carve out a distinct niche defined by the enemies they support, and by those they avoid (Hold 1977; Chesson & Kaung 2008). Specialized natural enemies can maintain species-rich plant communities by attacking their host plants where they are abundant, impeding their fitness relative to competitors that avoid the enemy (Janzen 1970; Connell 1971; Bever et al., 2015). Hence, large-scale gradients in biodiversity may be attributed to greater pressure from specialized herbivores and pathogens at lower elevations and latitudes with benign climates that are warmer, wetter, and less-seasonal (Schemske et al., 2009; Comita et al. 2014; Terborgh 2012; LaManna et al., 2017; Levi et al. 2019).

Recent advances in ecological metabolomics offer a promising approach to understanding complex biotic interactions and the chemical ecology of plant communities across biodiversity gradients (Sedio 2017; Sedio et al., 2021; Volf et al., 2023). Plant-chemical defenses (secondary metabolites) mediate biotic interactions and host-use relationships between plants and their natural enemies (Becerra 1997; Kursar et al., 2009; Salazar et al., 2016). In contrast to primary metabolites involved in the resource-acquisitive metabolism, secondary metabolites are organic molecules that mediate plant responses to abiotic or biotic stress and function as defenses against the ability of herbivores or pathogens to identify or digest the plant host. In response, herbivores and pathogens can evolve counters to plant chemical defenses, but often at the cost of generality (Ehrlich & Raven 1964; Schemske et al. 2009). Plant-enemy coevolution can result in host-use patterns that track plant secondary metabolites, promote chemical diversity and species richness in plant communities (Sedio & Ostling 2013), and mediate selection for chemical divergence among closely related plants (Becerra 1997; Kursar et al., 2009; Endara et al., 2017; Salazar et al. 2016). Plant-secondary metabolites have been shown to be more evolutionarily labile than other traits, as even closely related species can have very different metabolomes (Becerra 1997, Kursar et al. 2009). At evolutionary timescales this evolutionary lability is expected to result in less phylogenetic conservation of secondary metabolites among species found in warmer, wetter, more stable climates, where plant-enemy interactions are hypothesized to be stronger and more specialized (Wallace 1878; Dobzhansky 1950; Janzen 1970; Connell 1971).

Along elevational gradients, the abundance of herbivores and pathogens tends to decrease with elevation (Pellissier et al., 2014; Sam et al., 2020), while abiotic stress tends to increase with elevation. This can result in a tradeoff (Coley 1985) in which high-elevation plants may be expected to invest more in chemical defenses because compensatory regrowth of biomass lost to natural enemies is relatively more costly under unfavorable abiotic conditions and low nutrients (Defossez et al., 2018; Salgado et al. 2016). Furthermore, abiotic stress itself may select for investment in, and optimization of, specialized secondary metabolites that mediate plant stress response or protect against damage, such as from ultraviolet light (Volf et al., 2020). Yet unlike plant-enemy interactions that may undergo reciprocal coevolution, abiotic stress should select for convergence on shared, optimal traits (Asplund et al., 2022; Bakhtiari et al., 2021). On the other hand, high-elevation conditions may select for unique metabolites not found in lowland plants (Defossez et al., 2021). Perhaps because of such discordant selection, some studies have found non-linear, hump-shaped relationships between herbivory, plant secondary metabolite dissimilarity and elevational gradients (Sam et al., 2020; Volf et al., 2020).

Despite the importance of plant chemistry in mediating community dynamics, key gaps remain in our understanding of the chemical ecology of plant communities across elevational gradients. Few studies have simultaneously examined the roles of elevation, climate, species diversity, and phylogeny in shaping plant chemical variation along elevational gradients. A few studies have examined chemical variation over elevational gradients at the community scale in herbaceous grassland communities (Defossez et al., 2018; Defossez et al., 2021), but previous studies of woody plants have focused on single genera (Sam et al. 2020; Volf et al., 2020; Volf et al., 2023. In addition, the insight into the role that plant secondary metabolites play in generating biodiversity patterns has until recently been limited by their overwhelming diversity and the lack of untargeted approaches to study them at macroecological scales. Here, we overcome this obstacle by taking advantage of recent innovations in untargeted metabolomics based on mass spectrometry (Wang et al., 2016; Dührkop et al., 2019) that enable the study of chemical ecology at the scale of species-rich ecological communities such as tropical forests (Sedio 2017; Sedio et al., 2018).

In this study we explored the hypothesis that stronger selection by natural enemies at lower elevations shapes gradients in the diversity and evolution of plant secondary metabolites in tropical forests. We utilized data from 16 1-ha forest plots distributed across an ∼3000-m elevational-diversity gradient in the Bolivian Andes (17-137 tree species per plot). Using recent advances in large-scale chemical-metabolomic analytical techniques (Wang et al. 2016, Sedio 2017), we compared patterns of primary and secondary foliar metabolites in 473 tree species (906 unique species-plot combinations) to plot species diversity, elevation, climate, and phylogeny across the gradient to test four predictions: 1) Interspecific differences in plant-secondary metabolites will increase with tree species diversity; 2) Interspecific differences in plant-secondary metabolites will increase towards warmer, wetter, less-seasonal climates; 3) Plant species exhibit faster evolution of secondary metabolites (i.e., less phylogenetic signal) in high diversity communities; and 4) Plant species exhibit faster evolution of secondary metabolites (i.e., less phylogenetic signal) in warmer, wetter, and less-seasonal locations. Evidence in favor of these predictions would lend support to the hypothesis that variation in the strength of selection for interspecific divergence in secondary metabolites associated with climatic gradients contributes to the widespread elevational diversity gradient in trees.

## Methods

### Forest Plot Data: The Madidi Project

Floristic data were collected as part of the Madidi project (Jørgensen et al., 2015), a collaboration of more than two decades between the Herbario Nacional de Bolivia and the Missouri Botanical Garden to document the flora of the Madidi region in the Andes of Bolivia (Tello et al., 2015). The region ranges in elevation from lowland rainforests located at around 200 m above sea level (a.s.l.) to high mountains above 6,000 m a.s.l., above the tree line (Fuentes 2005, Bach & Grandstein 2011). The elevational gradient is covered by different forest types and encompasses a broad range of abiotic (climatic and environmental) conditions (Rafiqpoor & Ibish 2004; Friedman-Rudovsky 2012). The Madidi Project includes a total of 50 1-ha permanent plots ranging in elevation from 212 m to 3334 m above sea level. For this study, we selected a subset of 16 1-ha permanent plots in which leaves were sampled for chemical analyses and which spanned gradients in elevation (662-3324 m above sea level), climate, and tree-species richness (17-137 species per plot) (Table 1). The 16 plots include three seasonally dry, low-elevation forest plots, and 13 moist, montane forest plots (Table 1). Abundant genera include: *Miconia* (Melastomataceae)*, Sloanea* (Elaeocarpaceae), and *Ocotea* (Lauraceae) in the low elevation moist plots; *Weinmannia* (Cunoniaceae)*, Hedyosmum* (Chloranthaceae), and *Clethra* (Clethraceae) at high elevations (> 2500m); and *Weinmannia, Hedyosmum,* and *Clethra* in the seasonally dry plots. Tree species richness exhibits the typical negative relationship with elevation among the 13 moist forest plots, whereas the three seasonally dry forest plots exhibit a unique pattern of low species richness at low elevations. Within each plot, all woody plants (hereafter trees) with a diameter at breast height of at least 10 cm were spatially mapped, measured, and identified to a valid species or morphospecies. The majority of these plots have been censused a minimum of two times.

**Table 1.** Variation in tree species richness, elevation, annual precipitation (AP), and mean annual temperature (MAT) among 16 1-ha forest plots in the Madidi Project, Bolivia. The last column shows the number and percentage of species sampled for metabolomics analyses in each plot. Dry forest plots *italicized*.

| Plot Name | Species richness | Elevation (m) | AP (mm) | MAT (°C) | N trees sampled | N species metabolomes (% of all species) |
| --- | --- | --- | --- | --- | --- | --- |
| Chaqui 32 | 35 | 3116 | 975 | 11.9 | 85 | 22 (62%) |
| Fuerte 27 | 80 | 1900 | 1350 | 18 | 236 | 72 (90%) |
| Kanupa 44 | 17 | 3324 | 961 | 10.7 | 52 | 12 (70%) |
| Lomaka 40 | 137 | 1242 | 1332 | 21 | 248 | 109 (79%) |
| Lomasa 39 | 95 | 1054 | 1383 | 21.5 | 250 | 81 (85%) |
| *Pintat 5* | 48 | 880 | 1684 | 22 | 164 | 42 (87%) |
| *Resina 12* | 50 | 662 | 1858 | 23.1 | 141 | 45 (90%) |
| Sumpul 34 | 103 | 1223 | 1500 | 20.4 | 245 | 79 (76%) |
| Tapuri 45 | 43 | 2697 | 1013 | 15.2 | 121 | 35 (81%) |
| Tintay 24 | 117 | 1400 | 1358 | 19.7 | 283 | 87 (74%) |
| Tintay 25 | 94 | 1468 | 1357 | 20 | 253 | 73 (77%) |
| Titiri 42 | 34 | 2859 | 945 | 13.1 | 106 | 29 (85%) |
| Tocoaq 29 | 83 | 2407 | 1202 | 15.6 | 170 | 71 (85%) |
| Tocoaq 30 | 62 | 2510 | 1221 | 15.7 | 133 | 48 (77%) |
| Tocoaq 28 | 81 | 2200 | 1288 | 16.8 | 209 | 67 (82%) |
| *Yarimi 9* | 40 | 850 | 1698 | 22.2 | 129 | 33 (82%) |

**Table 2.**
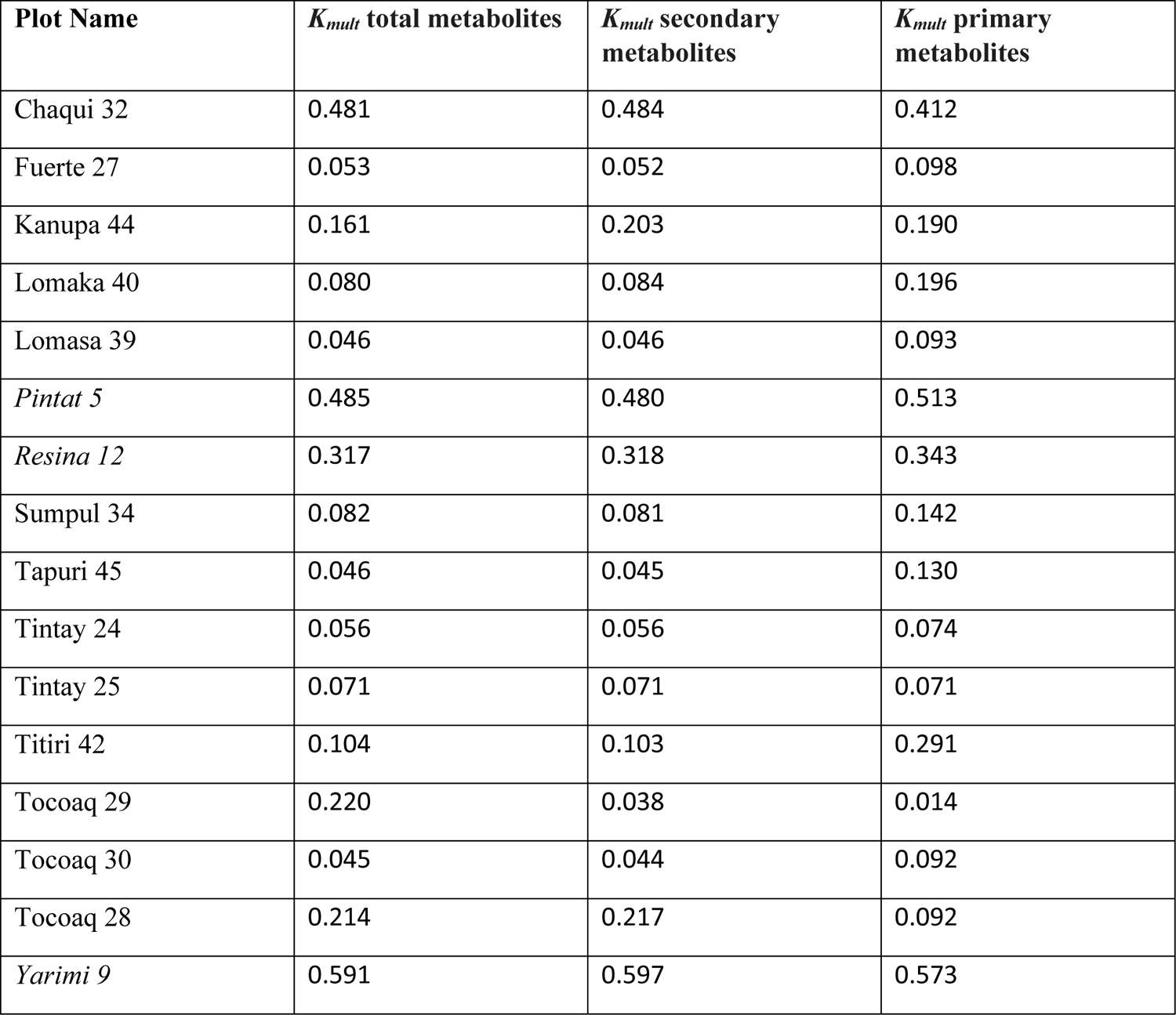
Phylogenetic signal (*K_mult_*) for all metabolites, secondary metabolites, and primary metabolites for each forest plot. Dry forest plots *italicized*.

| **Plot Name** | **$K_{mult}$ total metabolites** | **$K_{mult}$ secondary metabolites** | **$K_{mult}$ primary metabolites** |
| --- | --- | --- | --- |
| Chaqui 32 | 0.481 | 0.484 | 0.412 |
| Fuerte 27 | 0.053 | 0.052 | 0.098 |
| Kanupa 44 | 0.161 | 0.203 | 0.190 |
| Lomaka 40 | 0.080 | 0.084 | 0.196 |
| Lomasa 39 | 0.046 | 0.046 | 0.093 |
| *Pintat 5* | 0.485 | 0.480 | 0.513 |
| *Resina 12* | 0.317 | 0.318 | 0.343 |
| Sumpul 34 | 0.082 | 0.081 | 0.142 |
| Tapuri 45 | 0.046 | 0.045 | 0.130 |
| Tintay 24 | 0.056 | 0.056 | 0.074 |
| Tintay 25 | 0.071 | 0.071 | 0.071 |
| Titiri 42 | 0.104 | 0.103 | 0.291 |
| Tocoaq 29 | 0.220 | 0.038 | 0.014 |
| Tocoaq 30 | 0.045 | 0.044 | 0.092 |
| Tocoaq 28 | 0.214 | 0.217 | 0.092 |
| *Yarimi 9* | 0.591 | 0.597 | 0.573 |

### Chemical Analysis

#### Analytical chemistry and untargeted metabolomics

Across the 16 forest plots, we collected leaf samples from a total of 473 tree species, and 906 unique species-by-plot, for metabolomics analyses. Within forest plots, we collected leaf samples from 62–90% of the species in the plot (mean = 80% of species per plot; Table 1). Leaves of up to five individual trees per species per plot, that were collected after 2010, and dried on silica gel upon collection in the field were used for metabolite sampling. When a species had fewer than 5 individuals in a plot, we sampled leaves from all individuals. These samples were extracted for chemical analyses following Sedio et al. (2021). Briefly, 50 mg of dry leaf tissue was ground to a fine powder and 10 mg weighed for extraction in 1800 μL 90:10 methanol:water at pH 5 overnight at 4 °C. We used this solvent to extract metabolites over a wide range of polarity; mild acidity improves the solubility of most alkaloids, an important class of defensive secondary metabolites (Sedio et al., 2017). Extracts of five individuals per species per plot were pooled to create 906 extract pools representing each unique species-by-plot for subsequent analysis.

We analyzed filtered extract pools using ultra-high performance liquid chromatography-tandem mass spectrometry (UHPLC-MS/MS) using a Thermo Fisher Scientific (Waltham, MA, USA) Vanquish UHPLC with a C18 column and a Thermo QExactive quadrupole-orbitrap MS. Separation of metabolites by UHPLC was followed by heated electrospray ionization (HESI) in positive mode using full scan MS1 and data-dependent acquisition of MS2. Detailed instrumental methods are described by Sedio et al. (2021). Spectra for all 906 species-by-plot is curated as a public MassIVE dataset on the Global Natural Products Social (GNPS) Molecular Networking server (ftp://massive.ucsd.edu/MSV000090549)

Raw LC-MS data were centroided and processed for peak detection, peak alignment, and filtering using MZmine2 (Pluskal et al., 2010). Aligned chromatograms were used to create a ‘feature-based molecular network’ (FBMN; Nothias et al. 2020) using GNPS (Wang et al., 2016). The resulting network was used to create a dendrogram in which the structural similarities of all metabolites were reflected in one phylogeny-like dendrogram using Qemistree (Tripathi et al., 2021). Metabolites were annotated by predicting molecular formulae using Sirius (Dührkop et al., 2015), predicting molecular structures usig CSI:FingerID (Dührkop et al., 2019), and classifying compounds chemically using ClassyFire (Djoumbou Feunang et al., 2016) and NPClassifier (Kim et al., 2022).

NPClassifier is a deep-learning artificial intelligence that classifies metabolites according to basic biosynthetic pathways, in addition to superclasses and classes (Kim et al., 2022). We used the “pathway”-level classifications of NPClassifier to group metabolites into “primary” and “secondary” metabolites as follows: primary metabolites were those classified in the “Carbohydrates” and “Fatty acids” pathways, whereas secondary metabolites were those classified in the “Alkaloids”, “Amino acids and Peptides”, “Polyketides”, “Shikimates and Phenylpropoanoids”, and “Terpenoids” as well as compounds likely to be products of multiple core biosynthetic pathways that included one of these secondary-metabolite pathways. Glycosides were classified based on their non-carbohydrate moieties; nucleotides and nucleosides were classified among the carbohydrates at the “pathway” level (Kim et al., 2022). Our classification scheme was based on the broad likelihood of a metabolite being associated with anti-herbivore or antimicrobial defense. For example, amino acids and peptides include many primary metabolites, but may also include defensive compounds.

#### Chemical structural and compositional similarity (CSCS)

Sedio et al. (2017) developed a metric that quantifies chemical structural-compositional similarity (CSCS) over all compounds among species pairs. Conventional distance or similarity indices such as Bray-Curtis incorporate shared compounds but ignore structural similarity of unique compounds, and hence underestimate the similarity of species with distinct but very structurally similar, and perhaps functionally redundant, metabolites (Sedio et al., 2017). For each pair of the 906 species-by-plot, we calculated CSCS using i) the whole metabolome, using all metabolites in the data, ii) primary metabolites, and iii) secondary metabolites. We transformed CSCS matrices into dissimilarity matrices by calculating 1-CSCS. We calculated the abundance-weighted median 1-CSCS for the species assemblages represented by each of the 16 forest plots.

To disentangle the community chemical dissimilarity from the effect of diversity *per se*, we carried out rarefaction based on 12 species, the number of species sampled for chemical analyses in the most species-poor plot (Kañupa 44; Table 1). We calculated rarified CSCS by taking a random sample of 12 species and calculating the median chemical dissimilarity values at the plot-level. This operation was performed 1000 times for each plot and the mean of the distribution was taken as the rarified median chemical dissimilarity value. We obtained qualitatively similar results using observed and rarefied CSCS values. For simplicity, we therefore focus on results for observed CSCS and include results for rarefied CSCS in the supplementary material.

### Climate Data

Climate variables were selected to represent the variation in plot temperature, precipitation, and seasonality over the elevational gradient. Temperature variables included annual mean temperature and temperature annual range obtained from WorldClim Version 2.1 (Fick & Hijmans 2017). Precipitation variables included annual precipitation and precipitation seasonality, which is calculated as the ratio between the standard deviation and the mean precipitation of each month. The data were obtained from the Tropical Rainfall Measuring Mission (TRMM), a regional database that provides greater accuracy compared to WorldClim data in the Madidi region. The four variables were scaled and centered, and a principal components analysis was performed, of which the first two axes were used in the following analyses.

### Phylogenetic Signal

To quantify phylogenetic relationships among species, we constructed a phylogenetic tree using the V.Phylomaker package in R (Jin & Qian 2019). As inputs, V.Phylomaker requires family, genus and species information, which is then referenced against two combined mega trees (Zanne et al., 2014; Smith & Brown 2018) to generate the phylogenetic tree. The resulting tree was generated from all 50 of the Madidi permanent plots and had 1,123 unique species as tips. The tree was then rooted and transformed into a distance matrix using the ‘cophenetic’ function, in the stats package in R (RStudio Team 2020), in order to be directly comparable to the chemical distance matrices. The tree was pruned to include only the species recorded in the 16 plots for all phylogenetic signal analyses, which included 892 species-by-plot.

For each plot, we calculated Adams’ (2014) *K_mult_* metric of phylogenetic signal for multivariate trait data. This technique compares an Brownian-motion model of evolution in multivariate trait space to the observed trait data, accounting for the topology and branch lengths of the phylogeny. When *K_mult_* < 1, taxa are less chemically similar to one another than expected by Brownian motion evolution on the observed phylogeny, whereas *K_mult_* > 1 indicates that species are more chemically similar to each other than expected by Brownian motion. The *K_mult_* test is an improvement over the Mantel test for matrix correlation, which not consider an explicit model of trait evolution underlying the expected relationship between phylogenetic and trait distance (Swenson 2014).

### Hypothesis Testing: Plot-Level Regressions

We tested our predictions using linear regression. For each plot, we calculated tree species diversity as the inverse Simpson’s index using the R vegan package (Hurlbert 1971, Oksanen et al. 2017). We used the inverse Simpson’s index because it provides a scale-independent measure of diversity (effective number of species) that is insensitive to differences in numbers of individuals (Chase et al., 2018). To test Prediction 1, we regressed median chemical dissimilarity versus diversity (inverse Simpson’s index) for the 16 forest plots. To test Prediction 2, we regressed median chemical dissimilarity versus elevation and the first two principal component (PC) axes of climatic variation. To test Prediction 3, we regressed *K*_mult_ versus diversity, and to test Prediction 4, we regressed *K*_mult_ versus elevation and the first two PC axes of climatic variation among the 16 forest plots. The three seasonally dry, low-elevation forest plots appeared to exhibit distinct relationships to other variables not represented by the 13 moist, montane forest plots. Hence, all regressions were repeated with the 13 moist montane forests, excluding the seasonally dry forests.

## Results

### Overview of metabolomics and climate data

We detected 22,576 unique metabolites in foliar extracts from 473 tree species in 16 1-ha forest plots (Figure 1). Metabolites ranged in mass from 116.07 to 599.47 Daltons (Da). We generated a predicted molecular structure and biosynthetic classification for 18,364 and 19,844 of the metabolites, respectively. Metabolites classified at the level of biosynthetic pathway of origin (“pathway” in NPClassifier; Kim et al., 2022) were represented by 4,448 Alkaloids, 458 Amino acids and Peptides, 262 Carbohydrates, 928 Fatty acids, 584 Polyketides, 6,131 Shikimates and Phenylpropanoids, 6,871 Terpenoids, and 153 metabolites derived from more than one major pathway.

**Figure 1.**
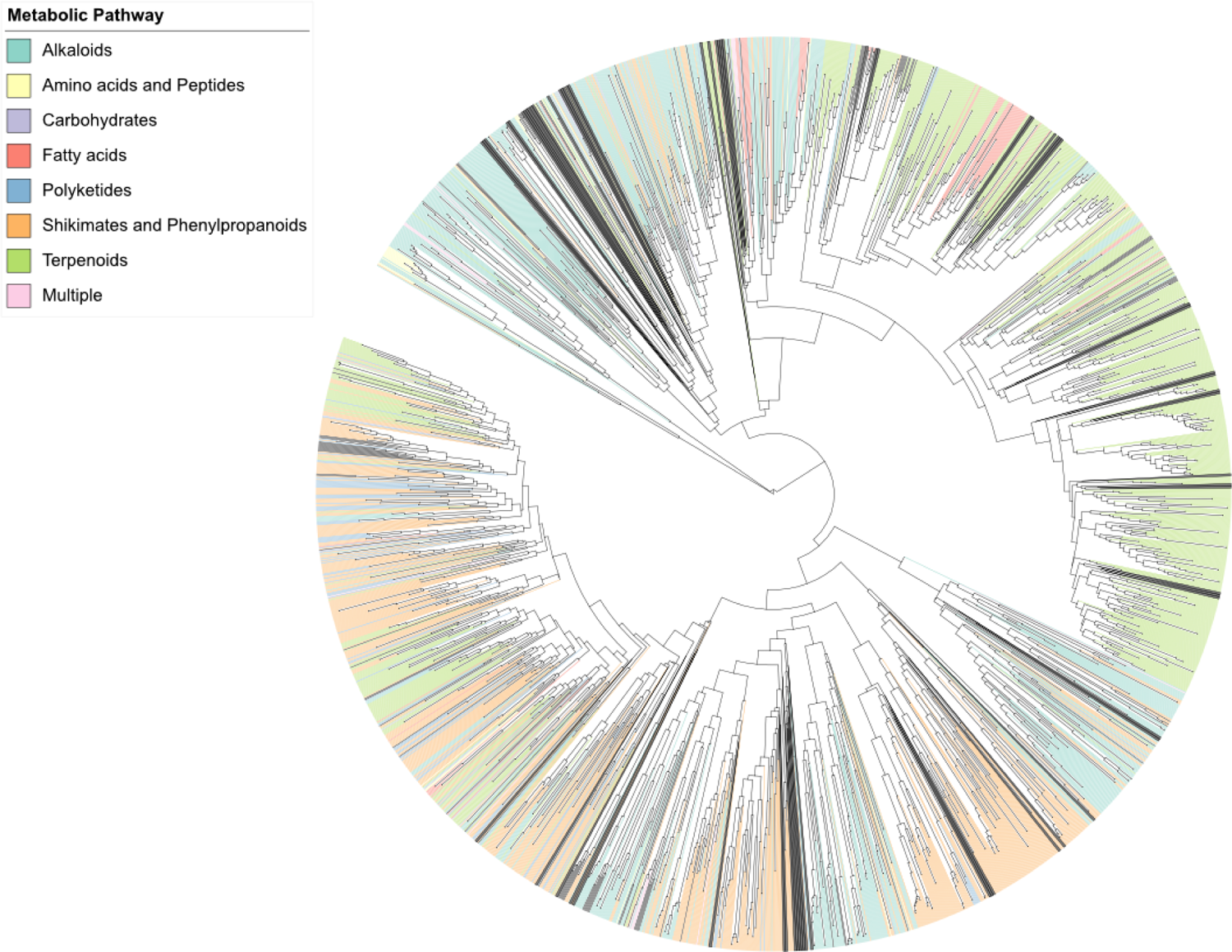
Qemistree dendrogram showing relationships of the 18,364 classified metabolites found within the 473 tree species sampled. Colors show primary metabolic pathways.

The first two principal components of climatic variation represented 71.2% and 15.8% of the variation among the 16 forest plots, respectively. The first principal component of climatic variation (PC1) primarily represents annual temperature and precipitation and annual temperature range, whereas the second principal component of climatic variation (PC2) primarily represents precipitation seasonality (Figure 2).

**Figure 2.**
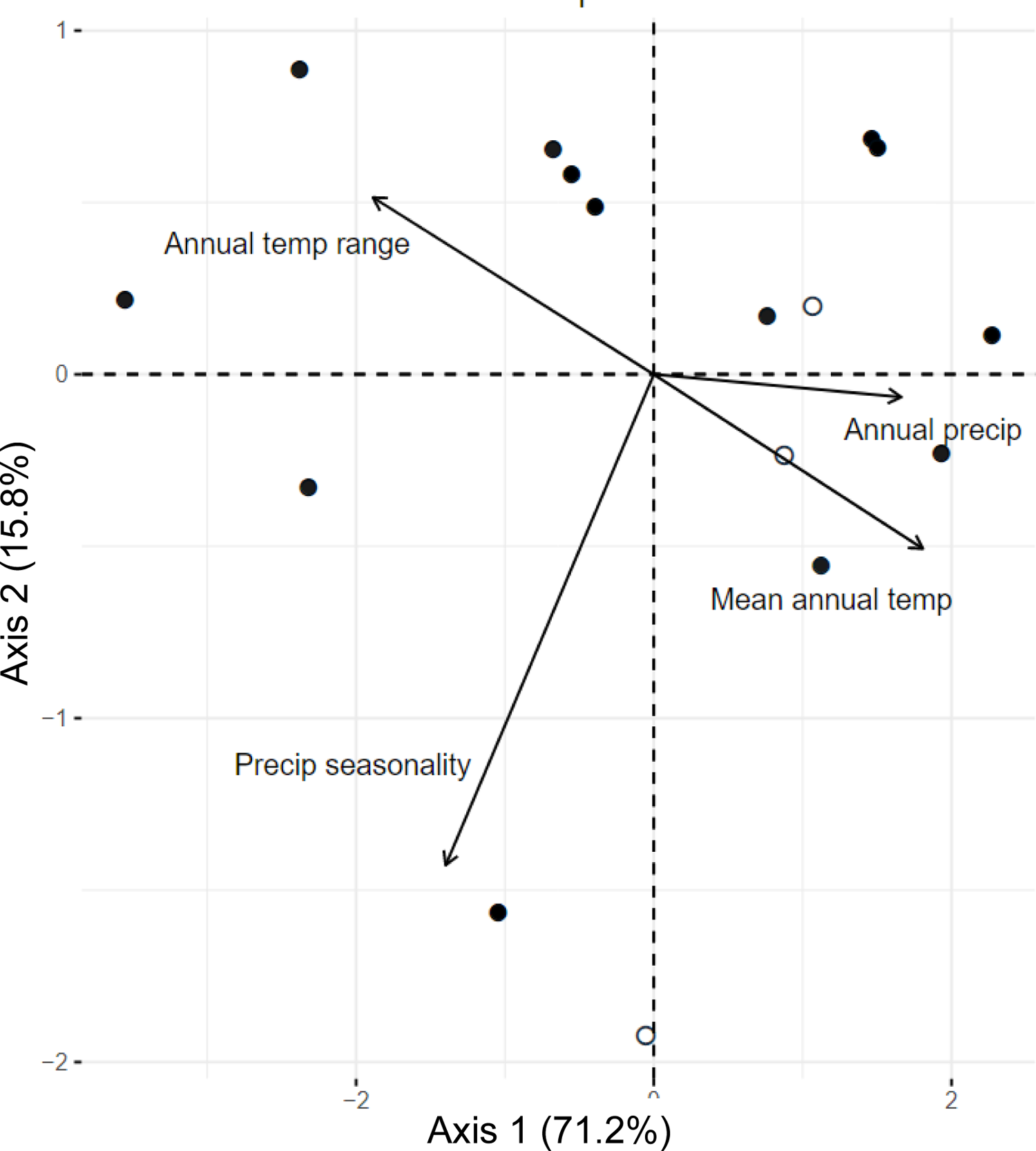
Principal Component Analysis (PCA) of four climate variables (mean annual temperature, mean annual temperature range, annual precipitation, precipitation seasonality) in the 16 forest plots. Arrows show the loadings of climate variable PC axis 1 and axis 2. Closed dots represent the moist forest plots, while open dots represent the seasonally dry forest plots.

### Chemical dissimilarity and phylogenetic signal across gradients

Chemical dissimilarity with respect to all detected metabolites increased with tree species diversity (R^2^ = 0.40, p < 0.01), decreased with elevation (R^2^ = 0.30, p = 0.02), and increased along Climate PC1 (R^2^ = 0.46, p < 0.01) but was unrelated to Climate PC2 among the 16 forest plots (Figure 3a-d). The relationship between chemical dissimilarity and elevation was much stronger when the three seasonally dry, low-elevation forest plots were excluded (Figure 3b, dashed line excluding open circles, R^2^ = 0.54, p < 0.01).

**Figure 3.**
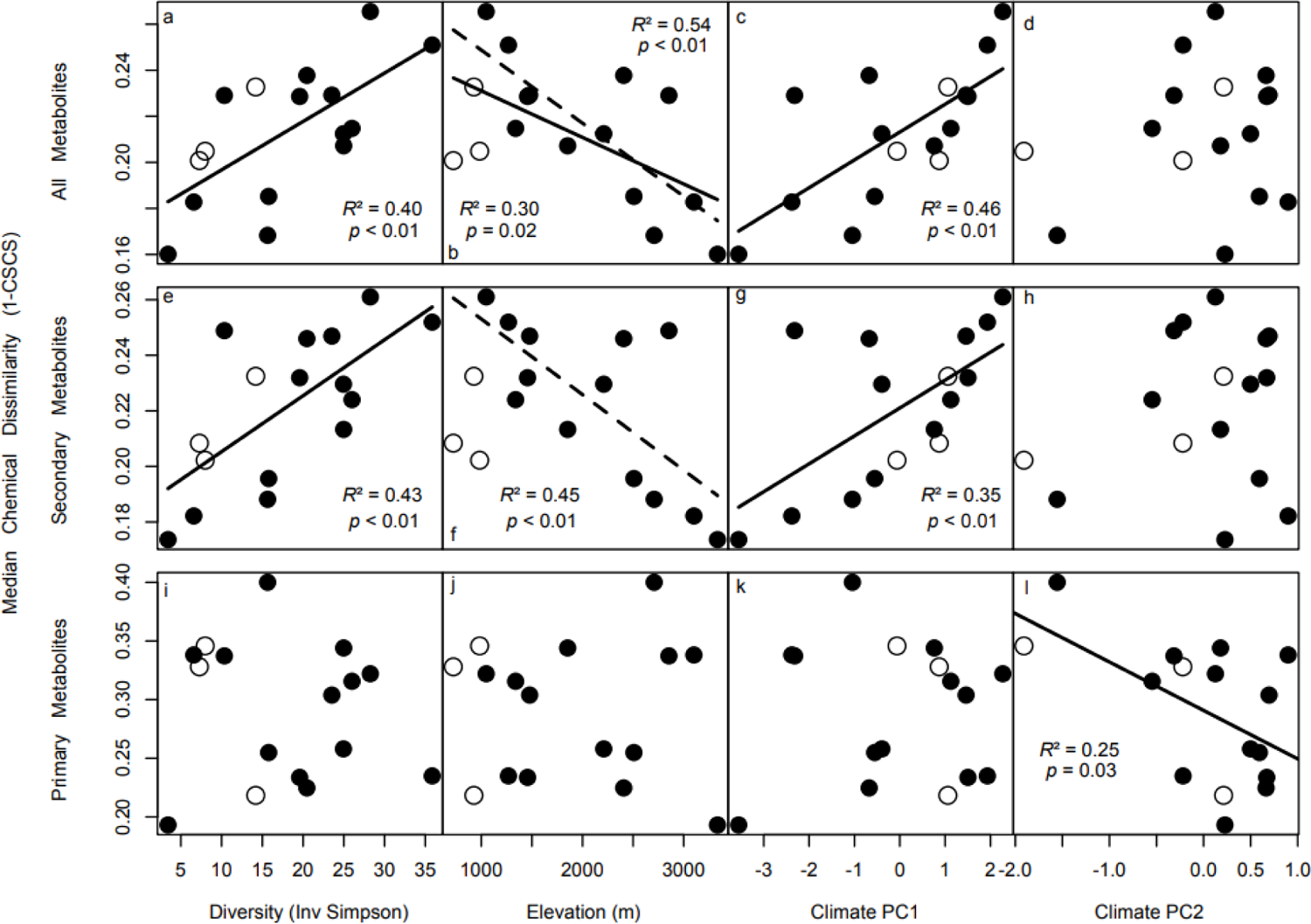
Relationships between median chemical dissimilarity (1-CSCS) and tree species diversity (inverse Simpson index), elevation (m), and climate among 16 forest plots in Madidi, Bolivia. Panels a-d represent linear regressions between median chemical dissimilarity among co-occurring species with respect to the whole metabolome and (a) species diversity, (b) elevation, (c) Climate PC1, and (d) Climate PC2, respectively. Panels e-h represent linear regressions between chemical dissimilarity with respect to secondary metabolites and (e) species diversity, (f) elevation, (g) Climate PC1, and (h) Climate PC2, respectively. Panels i-l represent linear regressions between chemical dissimilarity with respect to primary metabolites and (i) species diversity, (j) elevation, (k) Climate PC1, and (l) Climate PC2, respectively. Secondary metabolites are defined as those derived from the Alkaloids, Amino acid and Peptides, Polyketides, Shikimates and Phenylpropoanoids, and Terpenoids biosynethetic pathways using NPClassifier (Kim et al., 2022). Primary metabolites are defined as those derived from the Carbohydrates and Fatty acids pathways. Three seasonally dry forests (Table 1) are represented by open circles. Regressions using all 16 forest plots are represented by solid lines; regressions excluding three seasonally dry forests are represented by dashed lines. Adjusted *r*^2^ and *p*-values are presented for significant (*p* < 0.05) regressions only.

Chemical dissimilarity with respect to secondary metabolites increased with species diversity (Figure 3e; R^2^ = 0.43, p < 0.01) and Climate PC1 (Figure 3g; R^2^ = 0.35, p < 0.01) among the 16 forest plots. This measure decreased with elevation when the three lowland dry forests were excluded (Figure 3f; R^2^ = 0.45, p < 0.01). In contrast, plot median chemical dissimilarity with respect to primary metabolites decreased with climate PC2 (Figure 3l; R^2^ = 0.25, p = 0.03) but was unrelated to diversity, elevation, or climate PC1 (Figure 3i-k).

Phylogenetic signal was exceedingly low for all, secondary, and primary metabolites, as none of the plots approached the Brownian motion expectation (*K*_mult_ = 1) for any of the three metabolite classes (Table 1). Phylogenetic signal appeared greatest for low-elevation, low-species diversity dry forests and the highest-elevation, low-species diversity moist montane forests in the gradient (Table 1).

Phylogenetic signal with respect to all detected metabolites decreased with species diversity (Figure 4a; R^2^ = 0.27, p = 0.2) but was unrelated to elevation or Climate PC1 or PC2 (Figure 4b-d). Phylogenetic signal with respect to secondary metabolites decreased with species diversity (Figure 4e; R^2^ = 0.35, p = 0.01). When the three seasonally dry, low-elevation dry forest plots were excluded, phylogenetic signal with respect to secondary metabolites increased with elevation (Figure 4f; R^2^ = 0.26, p = 0.04) and decreased with Climate PC1 (Figure 4g; R^2^ = 0.24, p = 0.05). Phylogenetic signal with respect to primary metabolites decreased with species diversity (Figure 4i; R^2^ = 0.28, p = 0.02) but was unrelated to elevation or Climate PC1 or PC2 (Figure 4j-l).

**Figure 4.**
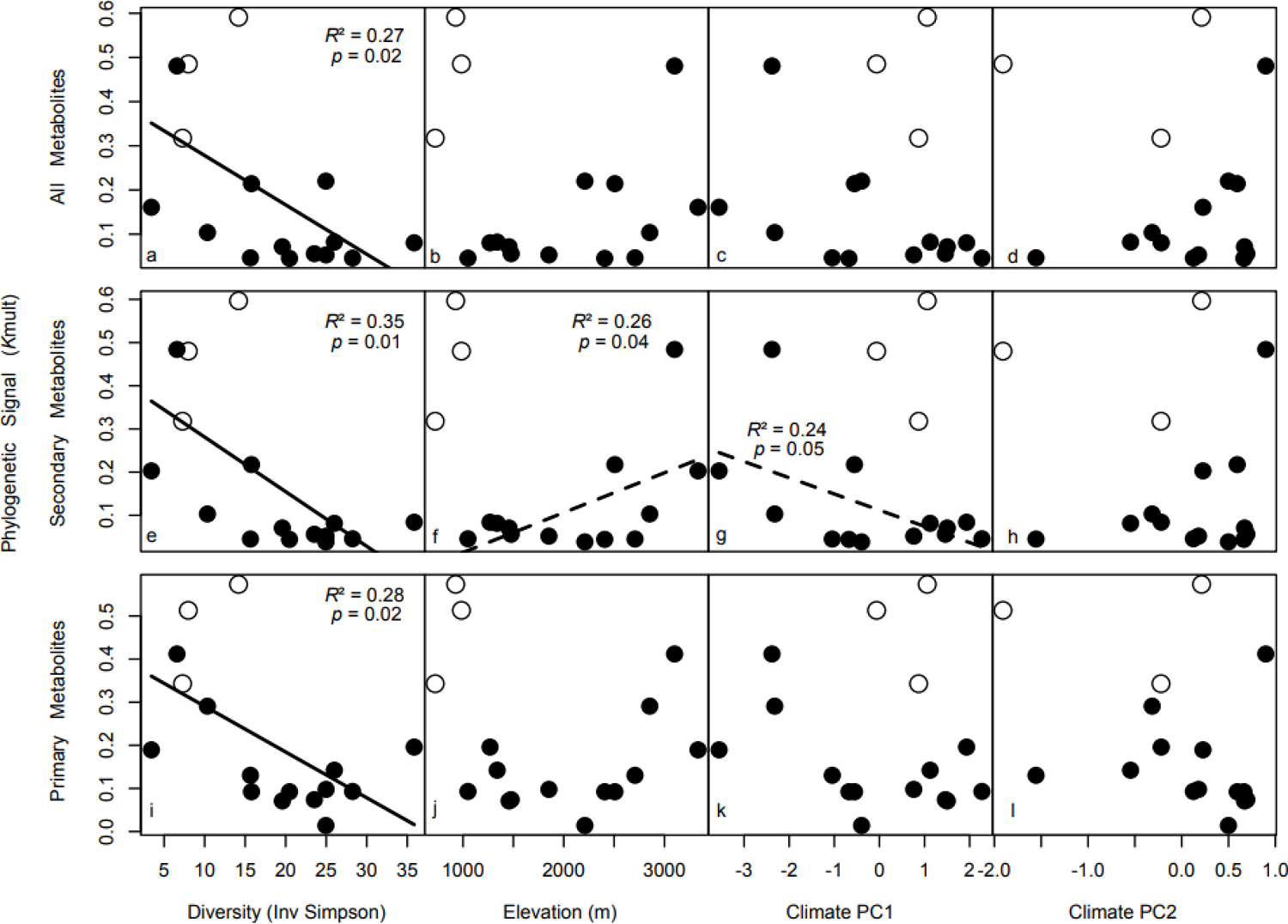
Relationships between chemical phylogenetic signal (*K*_mult_; Adams 2014) among co-occurring species and tree species diversity (inverse Simpson index), elevation (m), and climate among 16 forest plots in Madidi, Bolivia. Panels a-d represent linear regressions between phylogenetic signal among co-occurring species with respect to the whole metabolite and (a) species diversity, (b) elevation, (c) Climate PC1, and (d) Climate PC2, respectively. Panels e-h represent linear regressions between phylogenetic signal with respect to secondary metabolites and (e) species diversity, (f) elevation, (g) Climate PC1, and (h) Climate PC2, respectively. Panels i-l represent linear regressions between phylogenetic signal with respect to primary metabolites and (i) species diversity, (j) elevation, (k) Climate PC1, and (l) Climate PC2, respectively. Secondary metabolites are defined as those derived from the Alkaloids, Amino acid and Peptides, Polyketides, Shikimates and Phenylpropoanoids, and Terpenoids biosynthetic pathways using NPClassifier (Kim et al., 2022). Primary metabolites are defined as those derived from the Carbohydrates and Fatty acids pathways. Three seasonally dry forests (Table 1) are represented by open circles. Regressions using all 16 forest plots are represented by solid lines; regressions excluding three seasonally dry forests are represented by dashed lines. Adjusted *r* ^2^ and *p*-values are presented for significant (*p* < 0.05) regressions only.

## Discussion

Elevational diversity gradients are a striking feature of our planet and have inspired the development of ideas in ecology for centuries (von Humboldt & Bonpland 1807; Lomolino 2001; Rahbek 2005). The 16 tropical forest plots we examined here represented a wide range of variation in elevation, species diversity, and climate within a regional biodiversity hotspot in the central Andes Mountains. Classical hypotheses to explain large-scale diversity gradients posit that changes in species diversity are driven by geographic variation in the relative strength and nature of selection imposed by the abiotic and biotic environment (Wallace 1878; Dobzhansky 1950; Schemske et al. 2008; Lim et al., 2015). In turn, these processes are predicted to create systematic differences in the interspecific dissimilarity and phylogenetic signal of secondary-metabolite profiles among co-occurring tree species across ecological gradients. Our results broadly support four specific predictions concerning the relationships between chemical dissimilarity and phylogenetic signal and underlying gradients in species diversity, elevation, and climate, which we discuss below.

### High-diversity communities are composed of species with more dissimilar secondary metabolites

In Prediction 1, we predicted a positive relationship between chemical dissimilarity and community species diversity, based on the hypothesis that diversity itself is increased by antagonistic biotic interactions that select for chemical divergence among species (Dobzhansky 1950; Ehrlich & Raven 1964), reduce natural-enemy overlap among species (Becerra 1997; Endara et al., 2017), and promote competitive coexistence (Sedio & Ostling 2013). Our results supported Prediction 1. We found that chemical dissimilarity increased with species diversity (Figure 3a,e), whether measured in terms of the whole metabolome or metabolites classified as secondary metabolites based on their biosynthetic pathways of origin as predicted by NPClassifier (Kim et al., 2022). These patterns emerged in models based on median chemical dissimilarity of all co-occurring species at the forest-plot scale, and models based on rarified subsamples of co-occurring species that served to control for variation in species richness (Figure S1). These results are consistent with hypotheses that attribute community-scale variation in species diversity to variation in the strength of mechanisms that promote chemical divergence among species, such as pressure from relatively host-specific, but oligophagous herbivores and pathogens (Schemske et al. 2009; Lim et al., 2015). Furthermore, dissimilarity with respect to secondary metabolites strongly increased with species diversity (Figure 3e), while dissimilarity with respect to primary metabolites did not (Figure 3i). This contrast suggests that species differences in secondary metabolites, which include alkaloids, phenolics, polyketides, terpenoids (including steroid, or cardiac, glycosides), and non-protein amino acids that function as anti-herbivore and/or antimicrobial defenses, contribute to the diversity gradient.

### Communities in warmer, wetter, and less-seasonal climates are composed of species with more dissimilar secondary metabolites

In Prediction 2, we predicted that chemical dissimilarity of co-occurring species would decrease with elevation and increase in warmer, wetter, and less-seasonal climates, based on the hypothesis that chemically mediated plant-enemy coevolution that selects for chemical divergence among plants plays a greater role in these abiotically benign climates (Wallace 1878; Dobzhansky 1950). Our results supported Prediction 2. We found that chemical dissimilarity decreased with elevation and increased with temperature and precipitation as reflected in climatic PC1. For the metabolome considered as a whole, these patterns was true whether three lowland tropical dry forests were included or not (Figure 3b,c). For secondary metabolites, the positive relationship between chemical dissimilarity and PC1 was significant (Figure 3 g), whereas the negative relationship between chemical dissimilarity and elevation was significant only if three lowland tropical forests were excluded (Figure 3f).

Recent investigations of metabolomic variation associated with elevational gradients have considered single genera of woody plants, such as *Ficus* in Papua New Guinea (Volf et al., 2020) and *Salix* in Europe (Volf et al., 2023), and communities of herbaceous plants in Europe (Defossez et al., 2018; Defossez et al., 2021). However, none of these studies have examined chemical dissimilarity of co-occurring species at the level of entire tropical tree communities. Volf et al., (2023) found that low-elevation *Salix* were more dissimilar with respect to salicinoids, an important class of phenolic chemical defenses, than high-elevation willows, a result consistent with ours.

Previous studies have shown that co-occurring species are less chemically similar than expected by chance. This result has been reported for species-rich tree and shrub genera in the lowland Neotropics, including *Bursera* (Burseraceae) in Mexico (Becerra 2007), *Inga* (Fabaceae) in Panama and Peru (Kursar et al., 2009), *Piper* in Costa Rica (Salazar et al., 2016), and *Protium* (Burseraceae) in Peru (Vleminckx et al., 2018). Similar patterns have been found in *Ficus* in Papua New Guinea (Volf et al., 2018), Euphorbiaceae (principally *Macaranga*) in Yunnan, China (Huang et al., 2022) and in an assessment of seven species-rich genera in Panama (Sedio et al., 2017). In addition to studies focused on single lineages, community-wide studies have found that plants that co-occur within meters are more chemically dissimilar than expected from a community-wide sample (Henn et al., In Review; Yang et al. In Review) and the chemical dissimilarity of co-occurring species tends to decrease with latitude in the temperate and tropical zones (Sedio et al., 2018) and with temperature and precipitation within the boreal and temperate zones (Sedio et al., 2021).

Our study contrasts with these previous studies focused on single lineages and comparative studies focused on variation among whole plant communities at continental scales (Sedio et al., 2018; Sedio et al. 2021) in that we focused on variation along a local elevational gradient within the same biogeographic region (Central Andes). Nevertheless, our results are consistent with the widely observed chemical dissimilarity of tree species in low-elevation tropical forests. However, it is worth noting that Sedio et al. (2018, 2021) found differences in both the diversity of metabolites and in the chemical dissimilarity of species among geographically distant plant communities with very different biogeographic histories and little possibility for dispersal over ecological timescales. Our findings suggest that variation in temperature, precipitation, and seasonality over distances of kilometers may generate variation in community chemical dissimilarity comparable to that of climatic gradients on a continental scale, and hence that the underlying mechanisms that link climate to chemical evolution and competitive coexistence are likely general and operate over a wide range of variation in spatial scale (Zheng et al. 2021).

In stark contrast with our results concerning the whole metabolome or secondary metabolites, we observed a positive relationship between chemical dissimilarity with respect to primary metabolites and Climate PC2 (Figure 3l). As Climate PC2 was primarily defined by precipitation seasonality, this result suggests that species that occur in seasonal climates are more constrained in their chemical composition with respect to primary metabolites than species in less seasonal climates, which may be able to exploit a greater range of physiological variation associated with life-history strategy (Kambach et al. 2022).

### Chemical divergence among closely related species is greater in high-diversity communities, and in warmer, wetter, and less-seasonal climates

Phylogenetic signal was much lower than expected based on a model of Brownian motion without selection, even in high-elevation plots with comparatively higher phylogenetic signal than wetter, low-elevation plots (Table 1). Plants do exhibit phylogenetic signal with respect to broad chemical classes that tend to occur in certain plant families or genera (e.g., quinolizidine alkaloids in some lineages of legumes; Wink 2003). However, our result is consistent with other recent studies that have concluded that plant metabolite composition can be highly evolutionarily labile, especially in tropical climates when phylogenetic signal is measured among confamilial species (Becerra 1997; Kursar et al. 2009; Salazar et al. 2018; Volf et al., 2018; Huang et al., 2022), which can degrade phylogenetic signal when measured in the context of a species-rich forest community characterized by many co-occurring congeneric and confamilial species (Sedio et al., 2018; Sedio et al., 2021; Yang et al., In Review).

In Predictions 3 and 4, we predicted that chemical phylogenetic signal among co-occurring species would decrease with species diversity, increase with elevation, and decrease in warmer, wetter, and less-seasonal climates, based on the hypothesis that selection by natural enemies for chemical divergence among closely related species (Becerra 1997, Kursar et al. 2009) is relatively stronger in such environments. Our results supported Prediction 3 for secondary metabolites as well as both all and primary metabolites (Figure 4a,e,i). Prediction 4, that phylogenetic signal increases with elevation and decreases with temperature and precipitation reflected in Climate PC1, was supported only when we excluded the three seasonally dry, low-elevation forest plots (Figure 4 f,g). These and other results (Figure 3b,f) suggest that the climatic data we used in our PCA may not completely represent the climates experienced by the three seasonally dry forests. This may be because climate varies dramatically over a finer spatial scale in the topographically heterogeneous Andes and their foothills than that captured by the TRMM satellite, and/or the climatic variables we used did not reflect the source of abiotic stress experienced by the dry forests, such as, for example, if short periods of intense drought or strong interannual variation stresses plants in a manner that is not reflected in the TRMM precipitation seasonality variable. However, the relationships between secondary-metabolite phylogenetic signal and elevation and Climate PC1, respectively, changed when the three seasonally dry forests were excluded. This suggests that the rate of chemical divergence among tree species varies with temperature along a wet-forest elevational gradient. This result supports our Prediction 4, which follows from Dobzhansky’s (1950) hypothesis that biotic interactions are stronger forces of natural selection in warmer, wetter, and less-seasonal climates.

## Conclusions

Biodiversity gradients are a striking feature of our planet. While latitudinal biodiversity gradients have generally attracted more attention from biologists (Schemske et al., 2009), elevational gradients present perhaps a better opportunity to test hypotheses regarding proposed mechanisms (Graham et al., 2014), as key climatic variables vary over short distances, permitting experimentation or comparative study within a single system in which interacting species could plausibly disperse. Hence, we suggest that future research should take advantage of elevational gradients to test basic hypotheses concerning the intensity of plant interactions with insect herbivores and microbial pathogens, their effects on rates of chemical evolution in tree communities, and their contribution to the maintenance of species diversity.

Our results support the hypothesis that chemically mediated biotic interactions shape elevational diversity gradients by imposing stronger selection for interspecific divergence in plant chemical defenses in warmer, wetter, and less seasonal climates. Abiotic stress associated with high elevations may select for convergent secondary metabolite evolution distinct from that imposed by biotic stressors (Defossez et al., 2021; Volf et al., 2020; Volf et al., 2022), but competitive interactions among plants mediated by shared herbivores and pathogens are expected to select for chemical divergence. Our results suggest that the strength of this mechanism varies with climate in a manner that affects character evolution and diversity in plant communities. Our study also illustrates the promise of ecological metabolomics in the study of biogeography, community ecology, and complex species interactions in high-diversity ecosystems.

## Acknowledgements

We thank J. Dwenger for assistance in the lab, M. Volf for valuable discussion, and Adam Smith and Rachel Penczykowski for helpful comments on an earlier version of the manuscript. We thank the Dirección General de Biodiversidad, the Bolivian Park Service (SERNAP), the Madidi National Park and local communities for permits, access and collaboration in Bolivia, where fieldwork was supported by the National Science Foundation (DEB 0101775, DEB 0743457, DEB 1836353). Additional financial support to the Madidi Project was provided by the Missouri Botanical Garden, the National Geographic Society (NGS 7754-04 and NGS 8047-06), The German Embassy in Bolivia, International Center for Advanced Renewable Energy and Sustainability (I-CARES) at Washington University in St. Louis, the Comunidad de Madrid (Spain), Consejo Superior de Investigaciones Científicas (Spain), Centro de Estudios de América Latina (Banco Santander and Universidad Autónoma de Madrid, Spain), and the Taylor and Davidson families. Financial support for this study was provided by a grant from the Living Earth Collaborative at Washington University in St. Louis and the National Science Foundation (DEB Award 1557094 to JAM, DEB Award 2240431 to JAM, DEB Award 2240430 to BES). DH was supported by a Catharine M. Lieneman Botany Fellowship at Washington University in St. Louis.

