## Supplement for "Ecological metabolomics of tropical tree communities across an elevational gradient: Implications for chemically-mediated biotic interactions and species diversity"

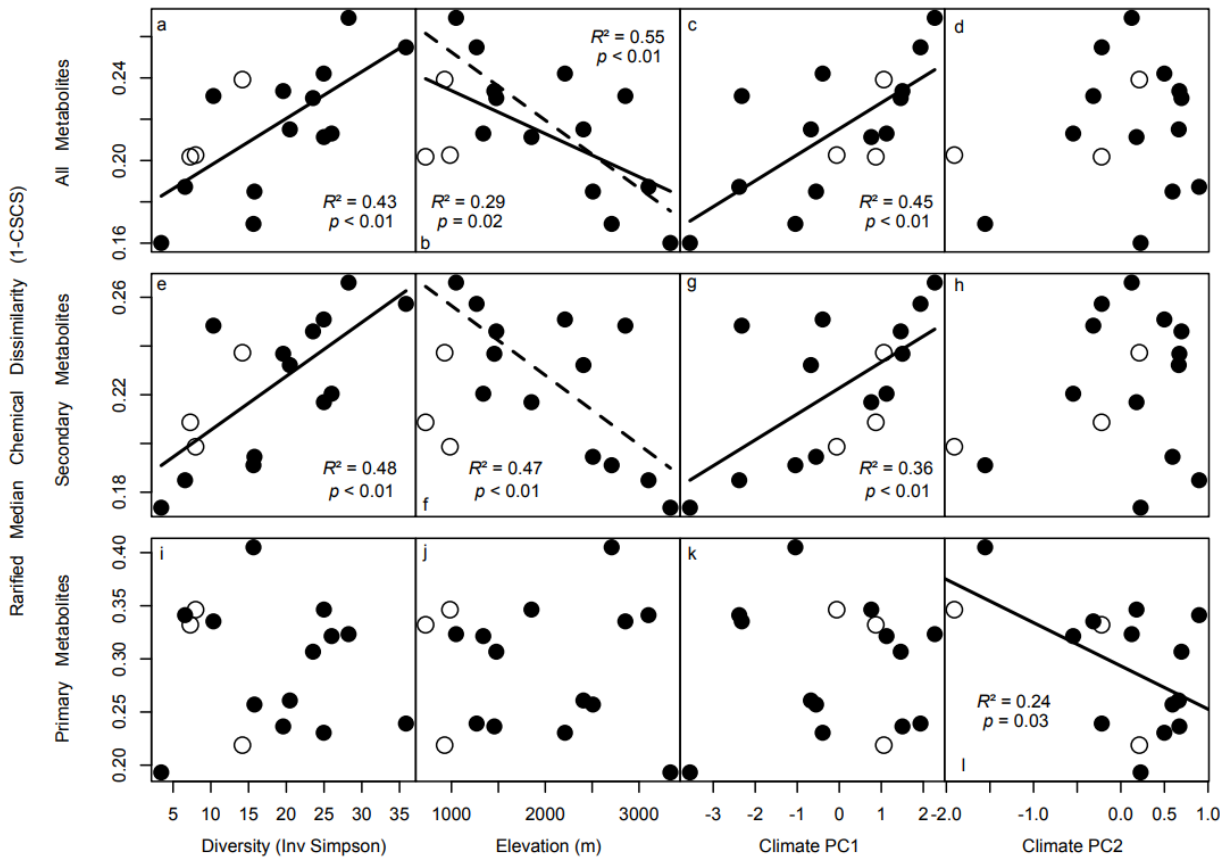


**Figure S1 –** Variation in rarified median chemical dissimilarity (1-CSCS) vs species diversity (inverse Simpson index), elevation (m), and climate among 16 forest plots in Madidi, Bolivia. Panels a-d represent linear regressions between rarified (n = 12) median chemical dissimilarity among co-occuring species with respect to the whole metabolite and (a) species diversity, (b) elevation, (c) Climate PC1, and (d) Climate PC2, respectively. Panels e-h represent linear regressions between chemical dissimilarity with respect to secondary metabolites and (e) species diversity, (f) elevation, (g) Climate PC1, and (h) Climate PC2, respectively. Panels i-l represent linear regressions between chemical dissimilarity with respect to primary metabolites and (i) species diversity, (j) elevation, (k) Climate PC1, and (l) Climate PC2, respectively. Secondary metabolites are defined as those derived from the Alkaloids, Amino acid and

Peptides, Polyketides, Shikimates and Phenylpropoanoids, and Terpenoids biosynethetic pathways using NPClassifier (Kim et al. 2022). Primary metabolites are defined as those derived from the Carbohydrates and Fatty acids pathways. Three seasonally dry forests are represented by open circles. Regressions using all 16 forest plots are represented by solid lines; regressions excluding three seasonally dry forests are represented by dashed lines. Adjusted $R^2$ and $p$-values are presented for significant ($p < 0.05$) regressions only.
